# Longitudinal clonal dynamics of HIV-1 latent reservoirs measured by combination quadruplex polymerase chain reaction and sequencing

**DOI:** 10.1101/2021.10.28.466316

**Authors:** Alice Cho, Christian Gaebler, Thiago Olveira, Victor Ramos, Marwa Saad, Julio Lorenzi, Ana Gazumyan, Susan Moir, Marina Caskey, Tae-Wook Chun, Michel Nussenzweig

## Abstract

HIV-1 infection produces a long-lived reservoir of latently infected CD4+ T cells that represents the major barrier to HIV-1 cure. The reservoir contains both intact and defective proviruses, but only the proviruses that are intact can re-initiate infection upon cessation of antiretroviral therapy (ART). Here we combine 4 color quantitative polymerase chain reaction and next-generation sequencing (Q4PCR) to distinguish intact and defective proviruses and measure reservoir content longitudinally in 12 infected individuals. Q4PCR differs from other PCR based methods in that the amplified proviruses are sequence verified as intact or defective. Samples were collected systematically over the course of up to 10 years beginning shortly after the initiation of ART. The size of the defective reservoir was relatively stable with minimal decay during the 10-year observation period. In contrast, the intact proviral reservoir decayed with estimated half-life of 4.9 years. Nevertheless, both intact and defective proviral reservoirs are dynamic. As a result, the fraction of intact proviruses found in expanded clones of CD4+ T cells increases overtime with a concomitant decrease in overall reservoir complexity. Thus, reservoir decay measurements by Q4PCR are quantitatively similar to viral outgrowth (VOA) and intact proviral DNA PCR (IPDA) with the addition of sequence information that distinguishes intact and defective proviruses and informs reservoir dynamics. The data is consistent with the notion that intact and defective proviruses are under distinct selective pressure, and that the intact proviral reservoir is progressively enriched in expanded clones of CD4+ T cells resulting in diminishing complexity over time.

**Significance:** HIV-1 infection requires lifelong treatment with antiretroviral therapy (ART) due to viral rebound of a latent reservoir of intact, transcriptionally silent provirus found to persist in the genome of CD4+ T cells. One of the major challenges to understanding the nature of the latent reservoir is accurately characterizing the measuring the size of the reservoir. Herein, we use quadruplex polymerase chain reaction (Q4PCR) to assess the dynamics of the latent reservoir in HIV+ individuals who have been on long-term ART for up to 10 years. Our results show that Q4PCR can be used to accurately measure the latent reservoir, while providing the added benefit of assessing the genetic diversity of the reservoir to better understand changes to clonal dynamics overtime.

## INTRODUCTION

HIV is a major global health problem, with an estimated 38 million people living with HIV leading to over 30 million HIV-related deaths as of 2020 (1, 2). While treatment with antiretroviral therapy (ART) effectively suppresses HIV-1 replication and reduces viremia (3), it does not cure infection. Treatment interruption typically leads to viral rebound within 2-4 weeks, emanating from a reservoir of intact transcriptionally silent proviruses that persist in the genome of CD4^+^ T cells (4–6). Thus, the intact proviral reservoir is the major barrier to HIV-1 cure (7–12).

Several different methods have been used to measure the HIV-1 latent reservoir and estimate its half-life many of which cannot distinguish between intact and defective proviruses (9, 13, 14). Interpreting the results of these measurements is complicated by the observation that the great majority of the proviruses integrated into the genome of CD4+ T cells are defective and that the decay rates for the two types of proviruses are different (13–19). For example, PCR measurements that rely on individual oligonucleotide probes do not distinguish between intact and defective proviruses and therefore any measure of HIV-1 decay by such methods is primarily a measure of the defective reservoir (20). Quantitative viral outgrowth assays (QVOAs) are limited to the intact proviral reservoir and underestimate the size of the reservoir because only a fraction of the latent cells in the cultures can be activated by stimulation *in vitro* (10, 21–23). Nevertheless, assuming that this fraction is stable, QVOA should be a relatively accurate measure of the rate of reservoir decay (21). The newly developed Intact Proviral DNA Assay (IPDA) is a high throughput assay that uses 2 sets of oligonucleotide probes to regions in the HIV genome that are conserved across most Clade B viruses (24). However, IPDA does not verify whether provirus sequences that hybridize to the 2 probes are fully intact (15, 24). As a result, IPDA may over-estimate the size of the reservoir and the accuracy of intact reservoir decay measurements may be variably altered by contaminating defective proviruses especially in individuals with small intact and large defective reservoirs (20). In addition, IPDA cannot be used to examine reservoir clonal dynamics. Q4PCR combines a 4-probe quantitative PCR strategy with near full-length amplification and sequence verification of intact or defective genomes (25). However, Q4PCR is a low throughput assay, and the requirement for full length genome amplification renders it less efficient than IPDA (20). Moreover, Q4PCR only captures the subset of defective proviruses that are detected by at least 2 of the 4 oligonucleotide probe sets (25).

To determine whether Q4PCR can be used to examine changes in the reservoir over time and document longitudinal reservoir clonal dynamics we sampled a cohort of 12 people living with HIV over a period of 10 years starting shortly after ART initiation. Reservoir half-lives measured by Q4PCR were consistent with those reported for QVOAs and IPDA during equivalent observation intervals (10, 11, 17, 19). As reported for IPDA (17), the decay rates of intact and defective proviral reservoirs differed significantly. Sequencing revealed that clonality increases and diversity decreases over time in both the intact and defective proviral reservoirs.

## RESULTS

### Characteristics of study participants

To investigate changes in the latent reservoir over time in people living with HIV on long-term ART, we assayed serum and peripheral blood mononuclear cell (PBMC) samples collected from 12 individuals who were treated with ART (Supplementary Table 1).

PBMC samples were collected at 3 or 4 time points: 1-9 months; 11-15 months; 2-3 years; and 5- or 10-years after initiating ART (Fig. 1A). For 3 individuals, additional plasma samples were collected 1-6 months before starting ART were used to compare circulating viral genomes to proviruses in the latent reservoir (Fig. 1A). All individuals had viral loads below 50 copies/mL at the evaluated time points, except for participant P7, who showed a minor viral blip at 59 copies/mL at the 2-3 year time point before returning to below 50.

**Figure 1.**
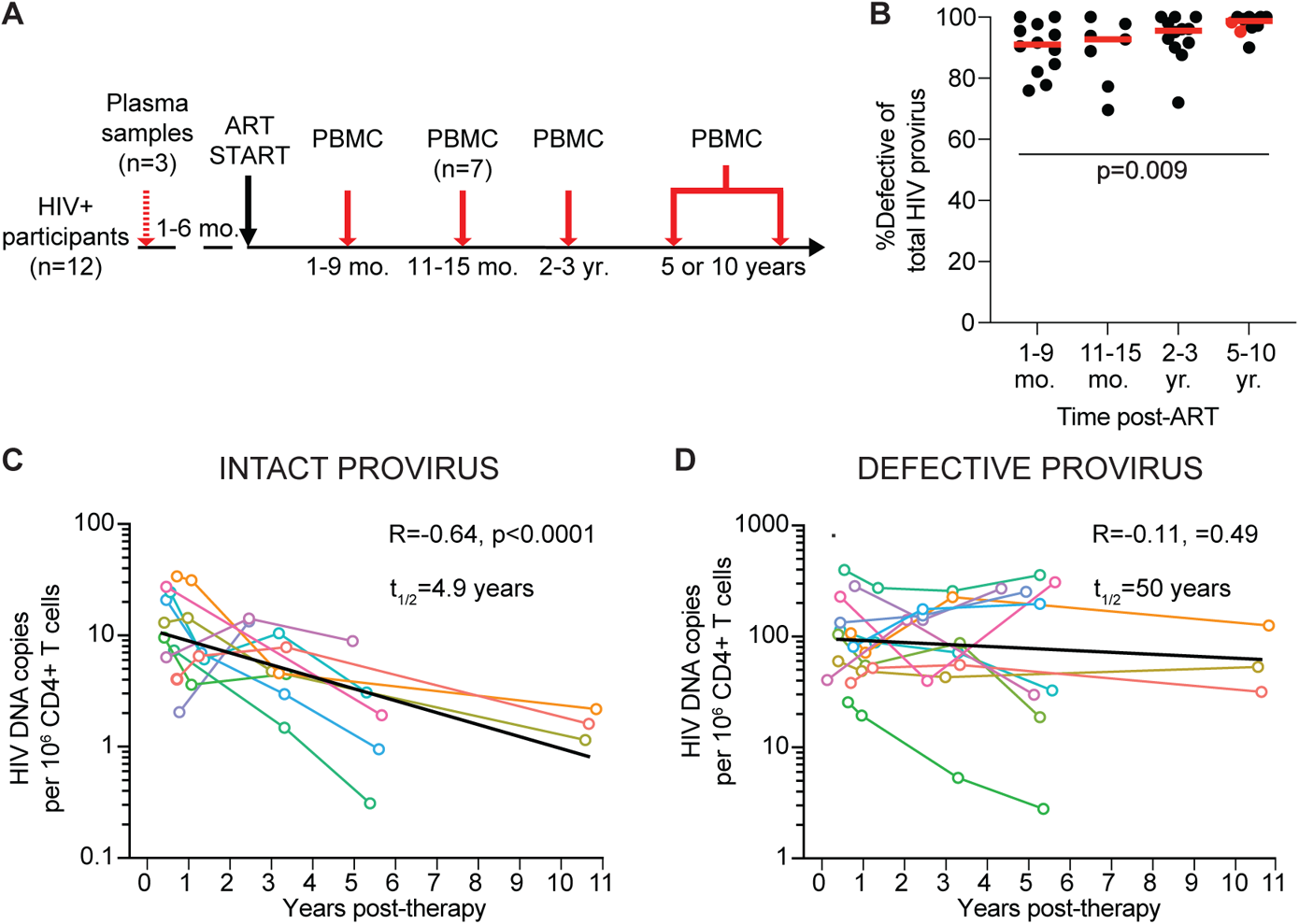
Longitudinal measure of the HIV-1 provirus latent reservoir by Q4PCR. (A) Study design. PBMC samples were collected from all 12 participants at 3-4 times points after starting ART, with plasma samples acquired from a subset (N=3) of participants prior to treatment. (B) Relative frequency of the defective HIV-1 proviral reservoir over time post-ART. Horizontal red line indicates median frequency. Participants sampled at 10 years post-ART are highlighted in red. (C) Number of intact HIV-1 proviruses per million CD4+ T cells detected overt time. Half-life of intact provirus reservoir was calculated to be 4.9 years (R=-0.64, p<0.0001). (D) Number of defective HIV-1 proviruses per million CD4+ T cells detected over time. The half-life of the defective provirus reservoir was measured to be 50 years (R=-0.11, p=0.49). Each colored line represents one individual participant. The exponential decay half-life was calculated by using log10 IUPM values and a linear regression model. Black line represents estimated decay of total data set. A Kruskal-Wallis test with subsequent Dunn’s multiple comparison’s test was used to analyze data where appropriate.

### Decay rate of intact and defective proviral reservoir

Sequence information was obtained for 323 intact and 4915 defective proviruses, with an average of over 100 sequences per participant per time point assayed (range: 9-355 sequences). On average 93% of the latent reservoir sampled by Q4PCR was comprised of defective proviruses at all time points assayed (Fig. 1B). For two participants, no intact sequences were detected at any time point, and an additional four participants had a single time point where no intact proviruses were detected (see Methods for additional information).

Individuals showed variable intact and defective reservoir dynamics (Supplementary Fig. 1). Nevertheless, when considered together there were significant differences between the two reservoir compartments. Whereas the intact proviral reservoir decayed with a half-life of 4.9 years (Fig. 1C, R=-0.64, p<0.0001), the defective proviral reservoir was far more stable with an estimated half-life of 50 years (Fig. 1D). As a result, there was a significant proportional increase in defective proviruses from 90% of the total HIV-1 reservoir at 1-9 months to 98% of sequences at the 5-10 year time point (p=0.009, Fig. 1B). We conclude that the half-life of the intact and defective proviral reservoirs in this cohort measured by Q4PCR differ, and that they are entirely consistent with the measurements reported by QVOA and IPDA in larger studies (11, 15, 17, 21).

### Reservoir evolution during long-term ART

To examine reservoir sequence evolution over time we compared circulating viral envelope (*env*) sequences obtained from plasma of 3 participants before they started ART with their corresponding proviral reservoir sequences from time points after ART initiation (Figure 1A). Although we did not find identical sequences between the two compartments, plasma and reservoir sequences clustered closely together (Fig. 2A).

**Figure 2.**
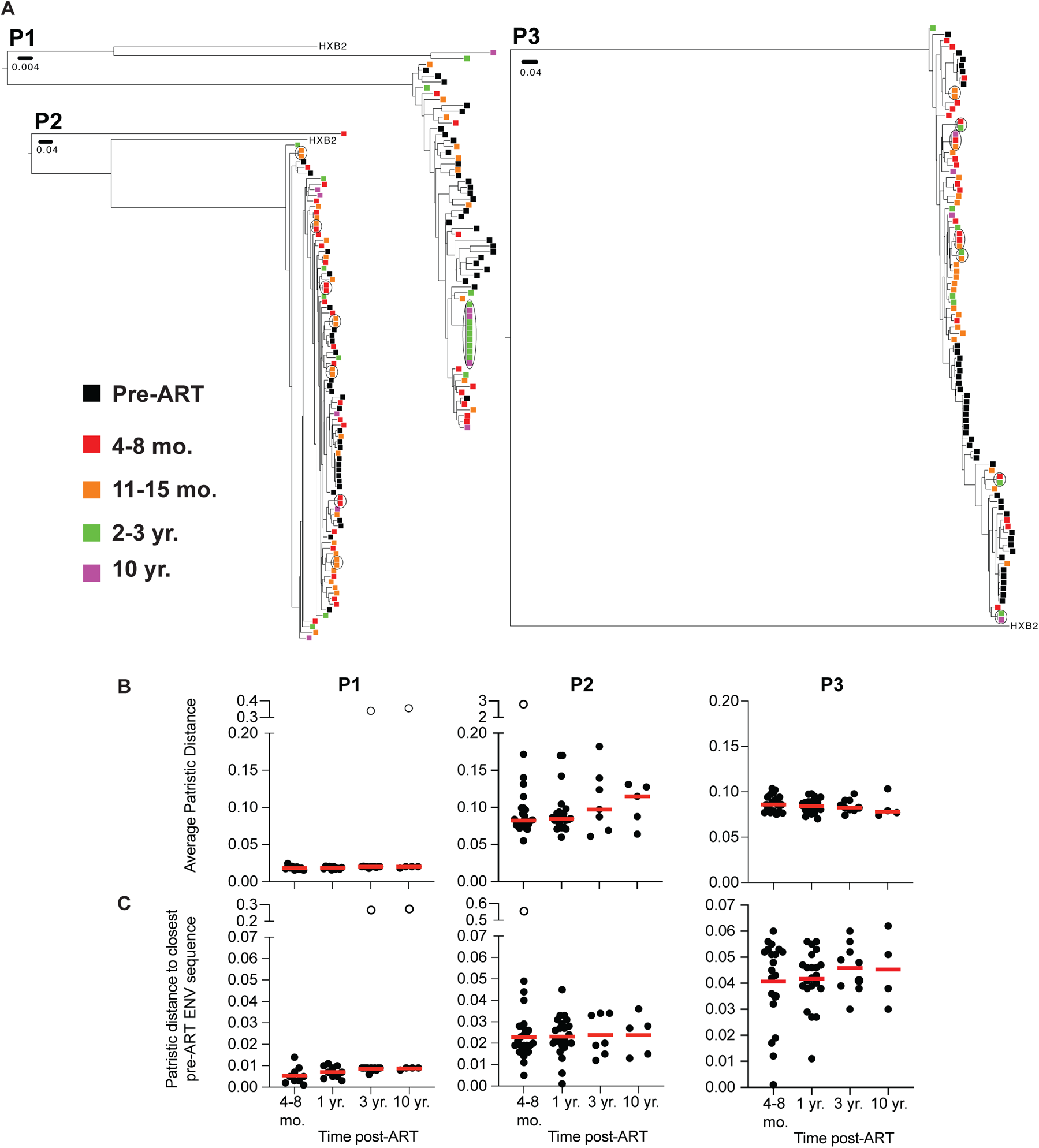
Provirus detected in circulation prior to ART are distinct from intact proviruses detected in latent reservoir found in PBMCs. (A) Phylogenetic tree of *env* sequences of intact proviruses in the latent reservoir and pre-ART proviruses found in circulation found in participants P1, P2, and P3. Branch lengths are proportional to a genetic distance (scale bar indicated below Sample ID). Black outlines indicate clones. Color of squares represent the origin of the sequence, as indicated by the key on the left. (B) Graph showing average patristic distance between each intact provirus found in the reservoir compared to every pre-ART virus found in circulation, comparing different time points post-ART. Each dot represents one intact provirus found in the reservoir. (C) Graph showing distance of intact proviruses found in the latent reservoir to the closest pre-ART *env* sequence. Each dot represents one intact provirus found in the reservoir. Participant name is specified above graphs. Unfilled dots indicate outliers that were excluded from the analysis. Horizontal red line indicates the median patristic distance at each time point. Grubb’s test was used to calculate outliers, and a Kruskal-Wallis test with subsequent Dunn’s multiple comparisons was used to analyze data where appropriate.

To further study the phylogenetic relationship between viruses circulating before ART initiation and proviruses in the reservoir we compared their average patristic distances. As reported by others (26), the average distance did not change overtime in any of the participants whether we considered all *env* sequences or only intact proviruses (Fig. 2B and Supplementary Fig. 2). Similar results were obtained when comparing the patristic distance between latent proviruses and their closest pre-ART genetic relatives (Fig. 2C). However, some of the intact proviruses in the early reservoir that were most closely related to circulating viruses disappeared confirming that many of the changes in the reservoir over time represent concomitant loss of variants and clonal proliferation, rather than diversification because of ongoing viral replication.

### Clonal dynamics in the defective proviral reservoir

Neither QVOA nor IPDA examine the genetic diversity of the reservoir (11, 17, 21, 24). As expected from the disproportionally greater number of defective proviruses, we retrieved more defective than intact proviruses at all time points: 1350 at 1-9 months; 728 sequences at 11-15 months; 1214 sequences after 2-3 years; 1623 sequences after 5-10 years (Fig. 3A). The lower number of sequences obtained at the 11-15 month time point reflects the smaller number of individuals sampled at that time point (Fig. 1A).

**Figure 3.**
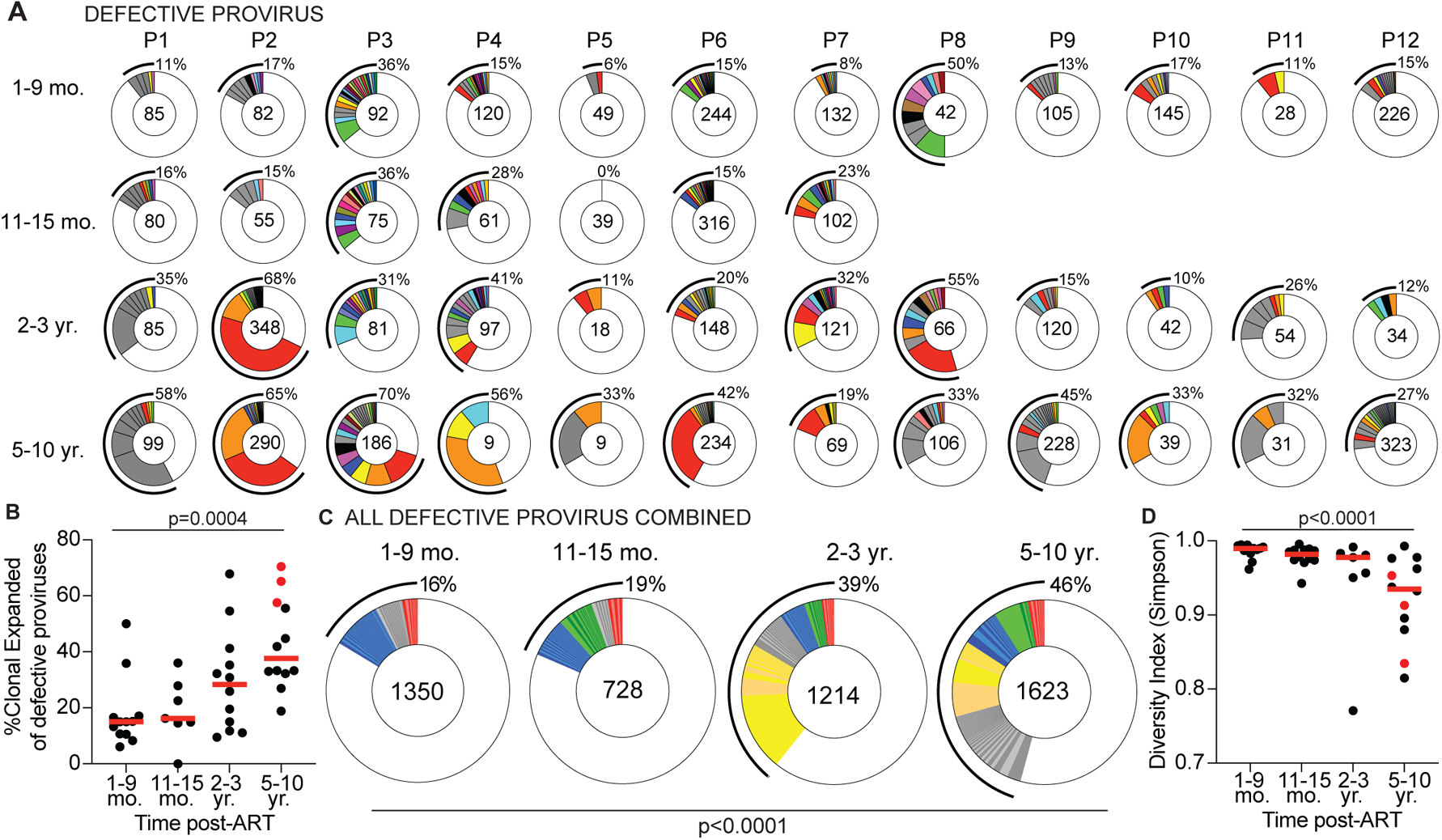
Clonal dynamics of defective provirus reservoir overtime. (A) Pie charts of the defective provirus reservoir detected in each individual, as denoted above the circles. Number inside each circle indicates the number of defective proviruses analyzed for each time point, specified to the left of the circles. Color slices indicate persisting defective clones found at multiple time points, while grey slices are clonal expansions unique to the time point. Remaining white slices represents sequences isolated only once. Black outlines indicate the frequency of clonally expanded defective proviruses detected in each participant. (B) Graphs showing frequency of clonally expanded defective provirus detected in participants over time. Each dot represents one individual. Participants sampled at 10 years post-ART are highlighted in red. Horizontal red bars indicate the median frequency of clonality at each time point. (C) Pie charts showing persisting or expanded clones of all defective proviruses from all individuals combined, organized by when the clone was first detected. Time point assayed is denoted above the circles. Number inside each circle indicates the number of defective proviruses analyzed for each time point. Color slices indicate the following: GREY – clonally expanded defective proviruses unique to the time point assays, BLUE – persisting clones of defective provirus, first detected at 1-9 months post-ART, GREEN – persisting clones first detected at 11-15 months post-ART, YELLOW – persisting clones first detected 2-3 year post-ART, RED – persisting clones detected at all assayed time points, WHITE – defective provirus sequences isolated only once. Black line indicates the frequency of clonally expanded defective proviruses detected at each time point. (D) Graph showing diversity (determined by Simpson’s Index) of all proviruses detected in participants over time. Each dot represents one individual. Participants sampled at 10 years post-ART are highlighted in red. Horizontal red bars indicate the median frequency of clonality at each time point. A Kruskal-Wallis test with subsequent Dunn’s multiple comparisons was used to analyze data where appropriate. A Fisher’s exact test with subsequent Bonferroni correction was used to compare clonality of defective proviruses between time points.

The overall representation of CD4+ T cell clones that contain integrated defective proviruses differs between individuals but increases over time (p=0.0004, Fig. 3B). When defective proviruses were considered in aggregate only 16% of all sequences were found in clones at the early time point and this increases to 46% after 5-10 years (p<0.0001, Fig. 3C). The defective reservoir was dynamic with only 3% of the clonally expanded proviruses persisting throughout the observation period (Fig. 3C). Clones found uniquely at one time point constitute 5%, 3%, 7%, and 17% of the overall reservoir at each of the 4 time points respectively (Fig. 3C). To determine how the increase in clonality may impact overall diversity, we used the Simpson Index to determine the probability that two randomly chosen proviruses are from the same clone (Fig. 3D). From this analysis, we see that the diversity significantly decreases over time (p<0.0001, Fig. 3D). Thus, the increased clonal representation results in a decrease of the overall proviral complexity in the reservoir.

The defective reservoir is composed of proviral genomes that contain undamaged open reading frames (ORFs) with the potential to encode HIV-1 proteins and others that cannot (27–30). For example, whereas proviruses with smaller internal deletions may be able to splice together mRNAs that encode some HIV-1 proteins, those with larger deletions, hypermutations, or mutations in the major splice donor (MSD) are less likely to be translation-competent and produce proteins (28, 31). It has been suggested that immune selective pressure, and therefore the kinetics of decay differs between defective proviruses that contain ORFs and can produce proteins and those that cannot (28, 31).

To determine whether the decay kinetics of different types of defective proviruses is heterogeneous, we separated defective proviral sequences into 5 categories: 1. Missing internal genes but able to encode protein products; 2. Non-coding that contain at least one early stop codon but possibly able to encode some functional proteins, 3. MSD mutation unable to produce spliced mRNA; 4. Inversions or Duplications; 5. Uncategorized defective proviruses (see Supplementary Table 2). All the different categories of defective proviruses show increased frequencies of clonally expanded sequences over time (Figure 4A-G, Supplementary Fig. 3).

**Figure 4.**
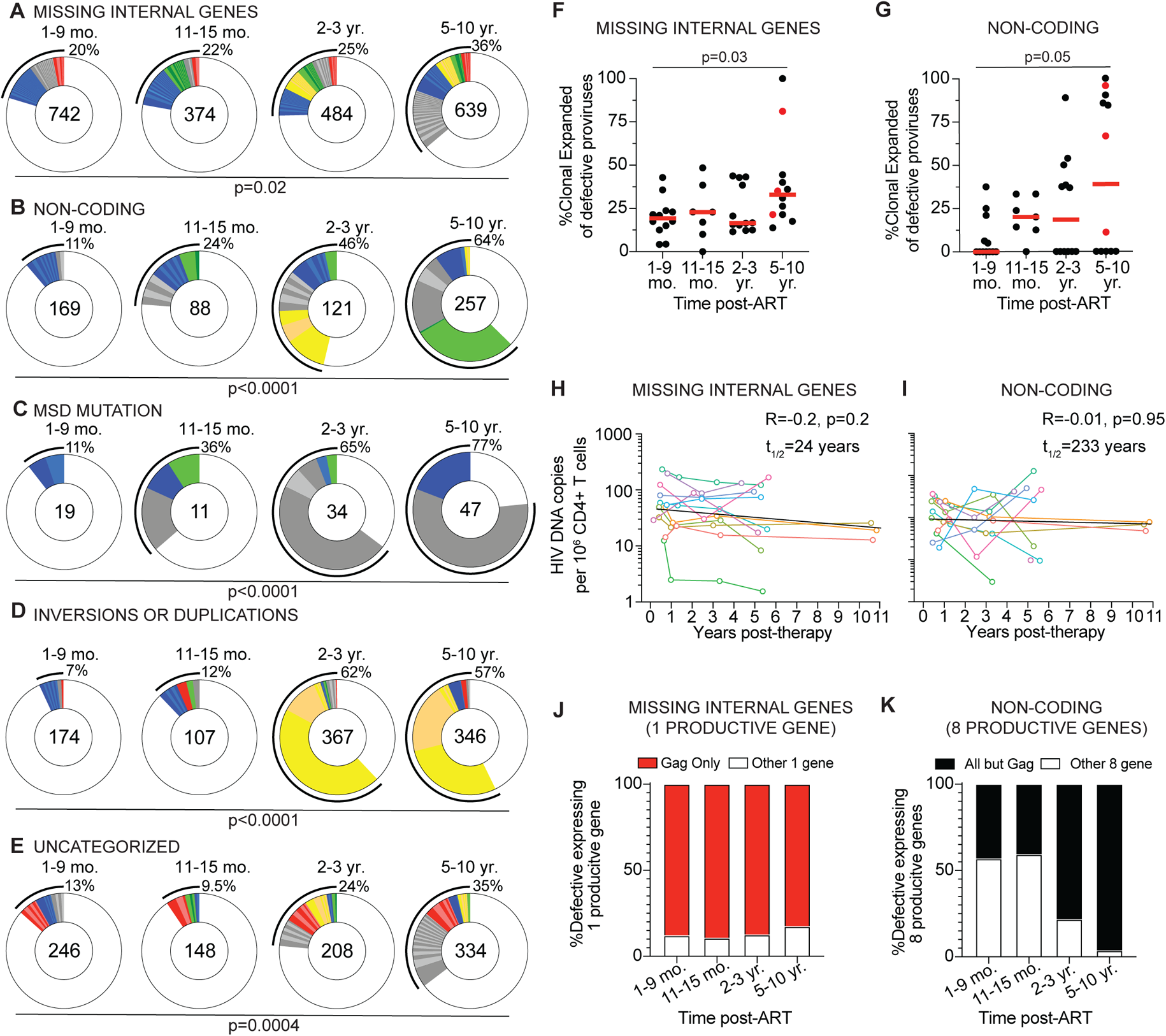
Longitudinal analysis of clonal dynamics of distinct types of defective proviruses. (A-E) Pie charts showing persisting or expanded clones of defective proviruses categorized as (A) missing internal genes, (B) non-coding, (C) MSD mutation, (D) Inversion or Duplications, and (E) Uncategorized. Time point assayed is denoted above the circles. Number inside each circle indicates the number of intact proviruses analyzed for each time point. Color slices indicate the following: GREY – clonally expanded defective proviruses unique to the time point assays, BLUE persisting clones of defective provirus, first detected at 1-9 months post-ART, GREEN – persisting clones first detected at 11-15 months post-ART, YELLOW – persisting clones first detected 2-3 year post-ART, RED – persisting clones detected at all assayed time points, WHITE defective provirus sequences isolated only once. Black outline indicates the frequency of clonally expanded defective proviruses detected at each time point. (F-G) Frequency of clonally expanded defective proviruses per individual over time, when looking at defectives categorized as (F) Missing Internal Genes and (G) Non-coding. Participants sampled at 10 years post-ART are highlighted in red. Red horizontal line indicated median frequency. (H-I) Graph showing the number of defective HIV-1 proviruses per million CD4+ T cells detected over time, when defectives are categorized as (H) Missing Internal (half-life=24 years, R=-0.2, p=0.2) and (I) Non-coding (half-life=233 years, R=-0.01, p=0.95). Each colored line represents one individual participant. (J) Graphs showing the frequency of Missing Internal Gene defective proviruses expressing 1 productive where the ORF expressed is *gag*. (K) Graph showing the frequency of Non-coding defective proviruses which are sequence predicted to express 8 functional proviral genes, where the ORFs expressed are all annotate genes excluding *gag.* A Kruskal-Wallis test with subsequent Dunn’s multiple comparisons was used to analyze data where appropriate. The exponential decay half-life was calculated by using log10 IUPM values and a linear regression model.

While overall clonality increased, there was variability in how specific clones wax and wane over time, with some defective proviruses showing large expansions at later time points (Figure 4A-E). One possibility is that these differences reflect ongoing responses to viral or other infections (32, 33). Consistent with the idea that defective proviruses that contain undamaged ORFs are under different selective pressure than those that are unlikely to be translation-competent, proviruses missing internal genes decayed faster than non-coding proviruses (Fig. 4H-I, Supplementary Fig. 3). The difference is likely due to *gag* expression because it is the dominant product of proviruses missing internal genes (Fig. 4J-K). As previously suggested (31), the disappearance of defectives that can produce *gag* protein could be mediated by cellular immunity.

### Clonal dynamics in the intact proviral reservoir

To examine clonal dynamics in the intact proviral reservoir we analyzed the intact HIV-1 proviral sequences from all participants across all time points. Overall, we obtained 152 intact proviral sequences from samples collected 1-9 months post-ART, 75 sequences from 11-15 months post-ART, 67 sequences from 2-3 years, and 29 sequences from 5-10 years post-ART (Fig. 5A). The decrease in the number of intact sequences detected from samples obtained at the 5-10 year time point is consistent with the decay kinetics of the intact latent reservoir (Fig. 1C).

**Figure 5.**
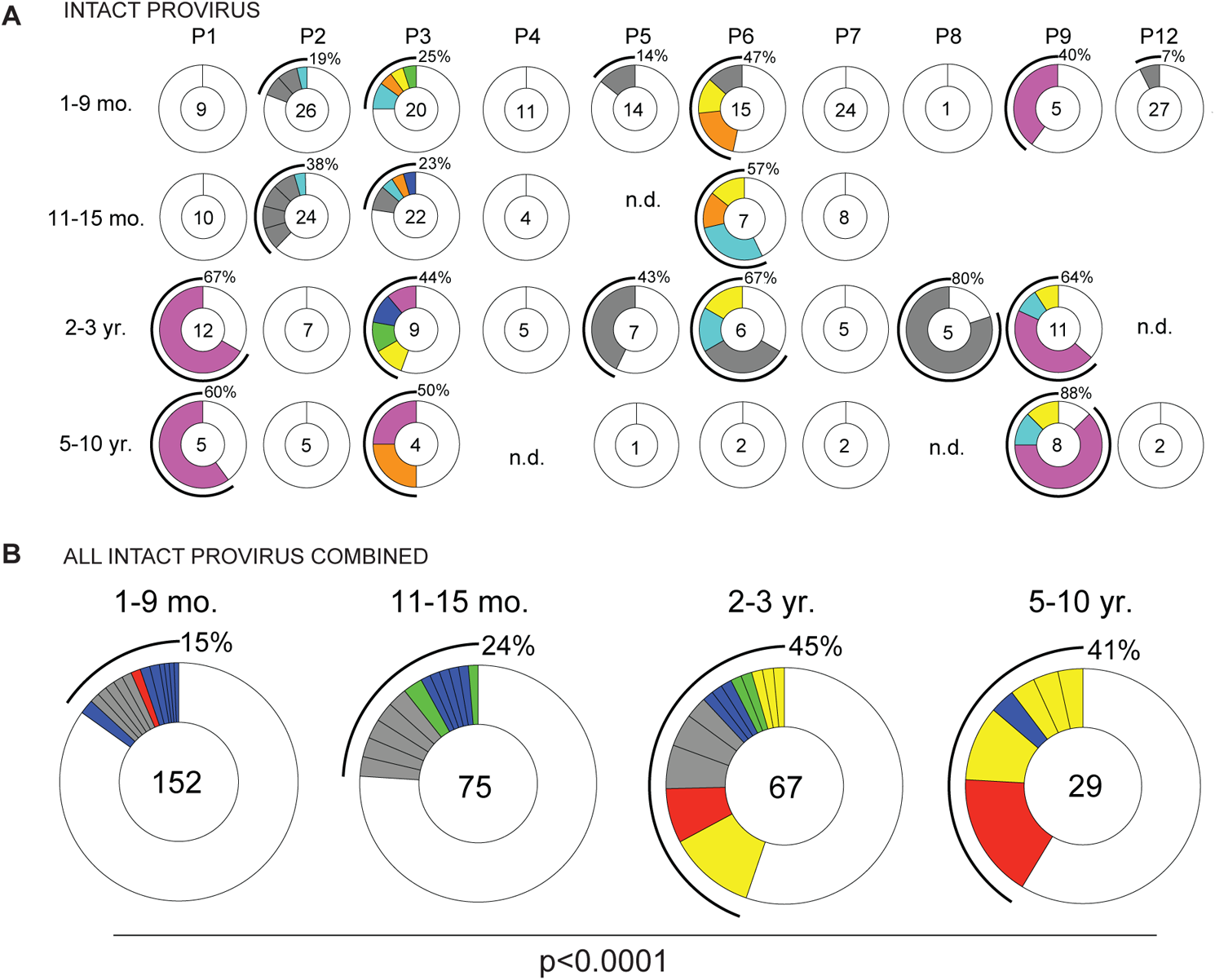
Clonal dynamics of the intact provirus reservoir overtime. (A) Pie charts of intact proviruses detected in each individual participant overtime. Participant name is denoted above the circles. Number inside each circle indicates the number of intact proviruses analyzed for each time point, specified to the left of the circles. Color slices indicate persisting intact clones found at multiple time points, while grey slices are clonal expansions unique to the time point. Remaining white slice represent sequences isolated only once. Black outlines indicate the frequency of clonally expanded intact proviruses detected in each participant. n.d. = not detected, no intact proviruses were detected from this sample. (B) Pie chart showing clonality of all intact proviruses detected in all individuals, combined, over time. Time point assays is denoted above the circles. Number inside each circle indicates the number of intact proviruses analyzed for each time point. Color slices indicate the following: GREY – clonally expanded intact proviruses unique to the time point assays, BLUE – persisting clones of intact provirus, first detected at 1-9 months post-ART, GREEN – persisting clones first detected at 11-15 months post-ART, YELLOW – persisting clones first detected 2-3 year post-ART, RED – persisting clones detected at all assayed time points, WHITE – intact provirus sequences isolated only once. Black line indicates the frequency of clonally expanded intact proviruses detected at each time point. A Fisher’s exact test with subsequent Bonferroni correction was used to compare clonality of intact proviruses between time points.

Expanded clones of CD4+ T cells containing intact integrated proviruses were present in all but 2 of the individuals examined (Fig. 5A). As expected from prior work (15, 23, 34–36) the intact proviral reservoir was dynamic with appearance and disappearance of some and persistence of other clones over time (Fig. 5A). When considered together the relative proportion of expanded clones in the latent reservoir increased with time from 15% at the first time point to 41% at the 5-10 year time point (p<0.0001, Fig. 5B). Thus, clonality increases in parallel with the decrease in size of the intact reservoir and as a result the complexity of the intact reservoir decreases with time. In conclusion, although the intact and defective reservoirs have different half-lives, they are both maintained in part by clonal expansion of CD4+ T cells.

## DISCUSSION

We performed a longitudinal analysis of the HIV-1 latent reservoir using Q4PCR to gain insights into the longevity and sequence evolution of the intact and defective reservoir over time. Although Q4PCR differs from other methods in a number of ways, the results were similar to QVOA and IPDA in the following respects: 1. The decay rate of the intact reservoir measured by Q4PCR (4.9 years) was comparable to that measured by VOA (mean: 4.4 years, range: 2.3-8.1 years) and IPDA (4-18.5 years) (9–11, 15, 17, 21); 2. Defective proviral decay was significantly slower than intact (15, 17, 31); 3. There was no evidence for proviral replication in the latent reservoir (26, 37); 4. Defective proviruses that have ORFs and can encode HIV-1 proteins decay faster than those that do not (27, 28, 31); 5. Increased defective proviral clonality overtime (15, 16, 38). In addition, the analysis revealed that the clonality of the intact latent reservoir increases and its diversity decreases over time.

Differences between Q4PCR and other methods could be due to any number of pitfalls associated with each of the assays. For example, VOA is labor intensive, variable from assay to assay (10), and ultimately underestimates the latent reservoir (22, 23). IPDA is a sensitive, high-throughput assay (24), but is limited to detecting proviruses that are sequence compatible with its 2 Packaging Signal (PS) and *env* oligonucleotide probe sets (18, 39). In addition, IPDA lacks a sequence verification step to determine if a provirus is truly intact or defective (20). The precision of the IPDA PS and *env* probes to identify genetically intact proviruses was estimated to be 70% in the original report (24). Similarly, a recently published direct comparison of IPDA and Q4PCR reported that of 625 probe positive samples identified by the PS and *env* probe combination, 305 (49%) were defective (20). This discrepancy is variable between individuals with some showing far larger or smaller differences than others (20). Nevertheless, the relative contribution of the more stable defective reservoir to the calculated IPDA half-life of the intact reservoir would be expected to increase over time as the intact reservoir decays. This potential problem may account for the increasing half-life of the intact reservoir measured by IPDA after 7 years and decrease the sensitivity of IPDA to changes in the intact reservoir over time (17).

Q4PCR is far more labor intensive than IPDA and not suited for the analysis of large clinical trials. In addition, the long-range PCR step in Q4PCR is less efficient that the short-range PCR used in IPDA, making Q4PCR a less sensitive assay that likely underestimates the size of the latent reservoir. We also limited our analysis to PBMCs isolated from blood samples. Although, blood appears to be representative of the overall latent reservoir (26, 40), it remains possible that we are missing information on reservoirs that reside solely in tissues. Despite these caveats, longitudinal studies show that half-life of the intact reservoir obtained by Q4PCR, VOA, the initial 7 years by IPDA is remarkably similar.

Sequence analysis of the defective reservoir by near full-length sequencing on 4 individuals revealed that defective proviruses that encode proteins have a shorter half-life than those that cannot (31). These protein products could trigger immune responses leading to inflammation or T cell exhaustion (27, 28). In addition, peptide products of these proteins presented on HLA may induce negative selection by cytotoxic CD8+ T lymphocytes (28). In contrast, translationally silent proviruses might evade the immune system and thereby remain more stable over time (31). Thus, Q4PCR analysis confirms and extends prior observations including the finding that the clonality of the defective reservoir increases over time (15, 16, 38).

It has been more challenging to examine the longitudinal clonal dynamics of the intact than the defective reservoir because of the relative difficulty in obtaining intact sequences at later time points. Q4PCR data reveals that clonality of the intact reservoir increases with time and as a result the reservoir becomes less diverse. The parallel increase in the proportion of proviral clones in the intact and defective reservoirs is consistent with the idea that clonal expansion of CD4+ T cell harboring latent proviruses is driven in part by antigenic stimulation (32, 33).

In conclusion, Q4PCR is a low throughput method that is complementary to VOA and IPDA. This new method may be especially valuable for analysis of small studies that aim to document how clinical interventions may alter latent reservoir dynamics.

## MATERIALS AND METHODS

### Study participants

Leukapheresis products were collected in accordance with protocol approved by the Institutional Review Boar of the National Institute of Allergy and Infectious Diseases, National Institutes of Health and registered on ClinicalTrials.gov, NCT00039689. All study participants provided written informed consent.

For two participants (P10 and P11), no intact proviruses were detected at the first time point (1-9 months post-ART) after assaying genomic DNA (gDNA) from at least 0.5 M (million) CD4+ T cells. For these participants, and average of 0.5 M CD4+ T cells (range: 0.15 M – 1 M) were assayed per time point. For all remaining participants (N=10), an average of 1.4 M (range: 0.5 M – 3.5 M) CD4+ T cells were assayed per time point, resulting in a limit of detection of 2 intact proviruses per million CD4+ T cells.

### CD4+ T cell isolation and genomic DNA extraction

Total CD4+ T cells were isolated from cryopreserved PBMCs using magnetic labeling and negative selection using a CD4+ T cell isolation kit (Miletnyi, #130-096-533). Genomic DNA (gDNA) was immediately isolated from 3-37 million purified C4+ T cells using phenol-chloroform (41). Briefly, CD4+ T cells were lysed in Proteinase K buffer (100 mM Tris, pH 8, 0.2% SDS, 200 mM NaCl, and 5 mM EDTA) containing 20 mg/mL Proteinase K (Invitrogen, #AM2548), incubated overnight at 55°C followed by gDNA extraction with phenol/chloroform/isoamyl alcohol extraction and ethanol participation, including an RNAse A step. A Qubit 3.0 Fluorometer and a Qubit dsDNA BR Assay Kit (Thermo Fisher Scientific, #Q32853) was used to measure DNA concentrations.

### Q4PCR

Q4PCR was performed as previous described (25). Briefly, an outer PCR reaction (NFL1) was performed on gDNA at a single-copy dilution, which was determined by a gag limiting dilution assay. Outer PCR primers include BLOuterF (5’-AAATCTCTAGCAGTGGCGCCCGAACAG-3’) and BLOuterR (5’-TGAGGGATCTCTAGTTACCAGAGTC-3’) (22). 1 μL of NFL1 PCR product was used as a template for Q4PCR reaction, using a combination of four probes that target conserved regions of the HIV genome. Each probe sets consist of a forward and reverse primer pair, as well as a fluorescently labeled internal hydrolysis primer/probe, as previously described (25): PS forward (5’-TCTCTCGACGCAGGACTC-3’), PS reverse (5’-TCTAGCCTCCGCTAGTCAAA-3’), PS probe (5’-/Cy5/TTTGGCGTA/TAO/CTCACCAGTCGCC-3’/IAbRQSp, Integrated DNA Technologies), ENV forward (5’-AGTGGTGCAGAGAGAAAAAAGAGC-3’); ENV reverse (5’-GTCTGGCCTGTACCGTCAGC-3’), ENV probe (5’-/VIC/CCTTGGGTTCTTGGGA-3’/MGB, Thermo Fisher Scientific), Gag forward (5’-ATGTTTTCAGCATTATCAGAAGGA-3’), Gag reverse (5’-TGCTTGATGTCCCCCCACT-3’), Gag probe (5’-/6-FAM/CCACCCCAC/ZEN/AAGATTTAAACACCATGCTAA-3’/IABkFQ, Integrated DNA Technologies), and Pol forward (5’-GCACTTTAAATTTTCCCATTAGTCCTA-3’), Pol reverse (5’-CAAATTTCTACTAATGCTTTTATTTTTTC-3’). Pol probe (5’-/NED/AAGCCAGGAATGGATGGCC-3’/MGB, Thermo Fisher Scientific). Thermostabe DNA polymerase was made in-house by transforming E. coli BL21 (DE3) pLys S cells with pOpen Taw plasmid (Gene and Cell Technologies) and induing the culture overnight with 1 mM IPTG. Cell pellet was lysed with lysozyme and the enzyme was extracted, as previously published (42), followed by purification using a heparin Sepharose 6HR column. After dialysis, Taq polymerase was mixed with anti-Taq tp7 antibodies (43) at a 1:10 ratio. Each Q4PCR was performed in a 10 μL final reaction volume containing 0.1 μL of polymerase in DNA polymerase PCR buffer supplemented with MgCl_2_ (Invitrogen 18067017), dNTP, Rox (Invitrogen #12223012), and containing the following primer and probe concentrations: PS forward and reverse primers at 21.6 μM with 6 μM of PS internal probe, *env* forward and reverse primers at 5.04 μM with 1.4 μM of *env* internal probe, *gag* forward and reverse primers at 2.7 μM with 0.75 μM *gag* internal problem, and lastly *pol* forward and reverse primers at 6.75 μM with 1.875 μM of *pol* internal probe. qPCR conditions were 94°C for 10 min, 40 cycles of 94°C for 15 sec, and 60°C for 60 sec. All qPCR reactions were performed in a 384-well plate format using the Applied Biosystem QuantStudio 6 or 7 Flex Real-Time PCR system. We use QuantStudio Real-Time PCR software version 2.2 (Thermo Fisher Scientific) for data analysis. The same baseline correction (start cycle 3; end cycle 10) and normalized reporter signal (ΔRn) threshold (0.2) was set manually for all targets/probes.

Fluorescent signal above the threshold was used to determine the threshold cycle. Samples with a threshold cycle value between 10-30 of any probe or prole combination were identified. Samples showing reactivity with two or more of the four qPCR probes were selected for a nested PCR reaction (NFL2). The NFL2 was performed using 1 μL of the NFL1 product as a template. Reactions were formed in 20 μL reaction volume using Platinum Taq High Fidelity polymerase (Invitrogen, #11-304-029) and PCR primers 275F (5’-ACAGGGACCTGAAAGCGAAAG-3’) and 280R (5’-CTAGTTACCAGAGTCACACAACAGACG-3’) (22) at a concentration of 800 nM.

### Library preparation and sequencing

All nested PCR products from NFL2 were subjected to library preparation. The Quit 3.0 Flourometer and Qubit dsDNA BR Assay Kit (Thermo Fisher Scientific, #Q32853) were used to measure DNA concentrations. Samples were diluted to a concentration of 10-20 ng/μL. Tagmentation reactions were performed using 1 μL of diluted DNA, 0.25 μL Nextera TDE1 Tagment DNA enzyme and 1.25 μL TD Tagment DNA buffer (Illumina, #20034198). Tagmented DNA was ligated to a unique i5/i7 barcoded primer combinations using the Illumina Nextera XT Index Kit v2 and KAPA HiFi HotStart ReadyMix (Roche, #07958927001), and then purified using AmPure Beads XP (Beckman Coulter, #A63881). 384 purified samples were pooled into one library and then subjected to paired-end sequencing using Illumina MiSeq Nano 300 V2 cycle Kits (Illumina, #MS-103-1001) at a concentration of 12 pM.

### HIV-1 sequence assembly and annotation

HIV-1 sequence assembly was performed by our in-house pipeline (Defective and Intact HIV Genome Assembler), which can reconstruct thousands of HIV genomes within hours via assembly of raw sequencing reads into annotated HIV genomes. The steps executed by the pipeline are described briefly as follows. First, we removed PCR amplification and performed error correction using clumpify.sh from BBtols and package v38.72 (https://sourceforge.net/projects/bbmap/). A quality control check was performed with Trim Galore package v0.6.4 (https://github.com/FelixKrueger/TrimGalore) to trim Illumina adapters and low-quality bases. We also used bbduk.sh from BBtools package to remove possible contaminant reads using HIV genome sequences, obtained from Los Alamos HIV database, as a positive control. After the overlapping reads are merged by BBMerge, we use a k-mer–based assembler, SPAdes v3.13.1, to reconstruct the HIV-1 sequences. The longest assembled contig is aligned using BLAST with HXB2 to set it in the forward orientation. Some of the contig’s nucleotides are potentially modified by aligning the high-quality HIV-derived reads. The modified contig is then annotated according to its alignment with HXB2 using BLAST. Sequences with double peaks, i.e., regions indicating the presence of two or more viruses in the sample (cutoff consensus identity for any residue <70%), or samples with a limited number of reads (empty wells ≤500 sequencing reads) were omitted from downstream analyses. In the end, sequences were classified as intact or defective, wherein intact sequences were determined by presence of 3’ LTR and 5’ PSI and lacking any fatal defects in the ORF of the 9 genes annotated by sequence analysis. Defective sequences were subdivided into more specific classifications according to their sequence structure: Major Splice Donor (MSD) Mutation, Non-coding, Missing Internal Genes, Uncategorized, or Inversions/Duplications.

### Clonal analysis

Clones were defined by aligning sequences of each classification (Intact, MSD Mutation, Non-coding, Missing Internal Genes, Uncategorized, or Inversions/Duplications) to HXB2 and calculating their Hamming distance. Sequences having a maximum of three differences between the first nucleotide of gag and last nucleotide of nef (reference: HXB2) were considered members of the same clone if found more than once across sequences from a single or multiple time points retrieved from each participant, based on the full amplification of a 9 KB template (99.667% identical).

### Single-genome analysis (SGA) of plasma virus *env* genes

Full-length *env* genes was amplified by nested PCR from plasma-derived viral cDNA, as previously described (44–46).Briefly, HIV-1 RNA was extracted from plasma samples using QIAamp Viral RNA Mini Kit (Qiagen, #52904). Followed by first-strand cDNA synthesis using Superscript III reverse transcriptase (Invitrogen, #18-080-044). 1 μL of cDNA diluted to single-genome per reaction was used as a template for first round *env* PCR, in a final reaction volume of 20 μL using Platinum Taq High Fidelity polymerase (Invitrogen, #11-304-029) and PCR primers envB5out (5’-TAGAGCCCTGGAAGCATCCAGGAAG-3’) and envB3out (5’-TTGCTACTTGTGATTGCTCCATGT-3’) at a concentration of 200 nM. Subsequently, a nested PCR using 1 μL of the first round PCR product in a similar PCR reaction using PCR primers envB5in (5’-CACCTTAGGCATCTCCTATGGCAGGAAGAAG-3’) and envB3in (5’-GTCTCGAGATACTGCTCCCACCC-3’). PCR conditions were 94°C for 2 min, 35 cycles of 94°C for 15 sec, 55°C for 30 sec, 68°C for 4 min, followed by a single cycle of 68°C for 15 min. Positive wells were determined using E-Gels (Invitrogen, #720841) and sequenced using Illumina MiSeq Nano 300 V2 cycle Kits (Illumina, #MS-103-1001) and analyzed, as described above. We assume all RNA recovered from plasma is reflective of intact viral particles. Sequence alignments, phylogenetic trees, and calculation of patristic distance to measure both the genetic distance and topology of the phylogenetic trees were performed by using Geneious Pro software, version 2020.0.3 and RAxML 8.2.11.

### Statistical analysis

Statistical analyses were performed using GraphPad Prism 9.

### Data Availability

All study data are included in the article and supporting information.

## Acknowledgements

We thank all study participants who devoted time to our research, and all members of the M.C. Nussenzweig laboratory of helpful discussions and Maša Jankovic for laboratory support. This work was supported by the Bill and Melinda Gates Foundation (Collaboration for AID Vaccine discovery grants OPOPP1092074, OPP1124068, and OPP1168933) and the National Institutes of Health (grants 1UM1 AI100663 and R01AI129795) to M.C. Nussenzweig; the Einstein– Rockefeller–CUNY Center for AIDS Research (grant 1P30AI124414-01A1); REACH-HIV Delaney (grant UM1 AI164565 to M. Caskey); and the Robertson Fund. T.W. Chun and S. Moir were supported in part by the Intramural Research Program of the National Institute of Allergy and Infectious Diseases, National Institute of Health. C. Gaebler was supported by the Robert S. Wennett Post-Doctoral Fellowship, and in part by the National Center for Advanced Translational Sciences (National Institutes of Health Clinical and Translational Science Award program, grant UL1 TR001866), and by the Shapiro-Silverberg Fund for the Advancement of Translational Research. M.C. Nussenzweig is a Howard Hughes Medical Institute Investigator.

The authors declare no competing interest. Correspondence may be addressed to

## Author contributions

C. Gaebler, A. Cho, M.Caskey and M.C. Nussenzweig designed the research. A. Cho, M. Saad, J.C.C. Lorenzi carried out experiments. A.G. produced reagents. A. Cho, C. Gaebler, T. Olveira, V. Ramos, M. Caskey and M.C. Nussenzweig analyzed the data. S. Moir and T.W. Chun recruited patients and executed clinical protocols. A. Cho and M.C. Nussenzweig wrote the manuscript.

## FIGURE LEGENDS

**Table S1.**
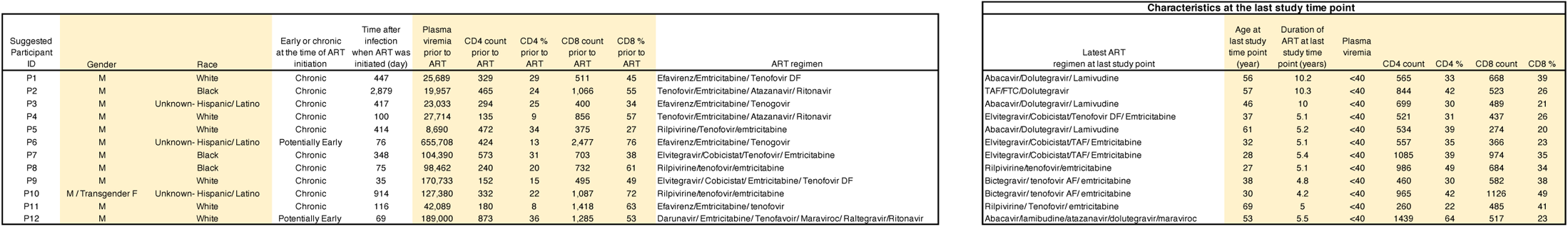
Characteristics of study participants.

**Supplementary Figure 1.**
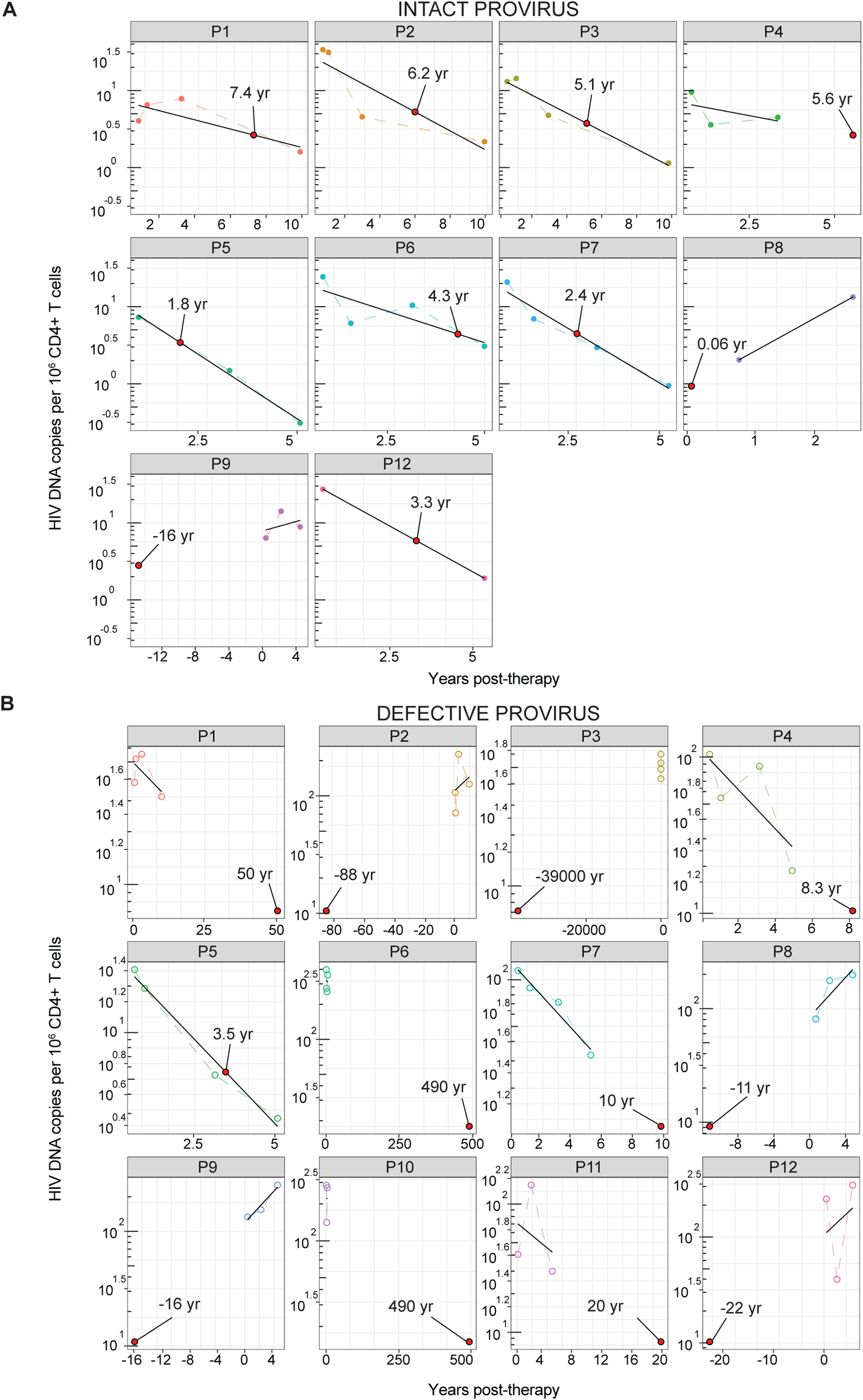
Half-life of the reservoir in each individual participant. The half-life of HIV proviral reservoir was calculated for each individual participant, for both (A) intact and (B) defective proviruses. Half-life is indicated by labeled red dot on each graph. The exponential decay half-life was calculated by using log10 IUPM values and a linear regression model.

**Supplementary Figure 2.**
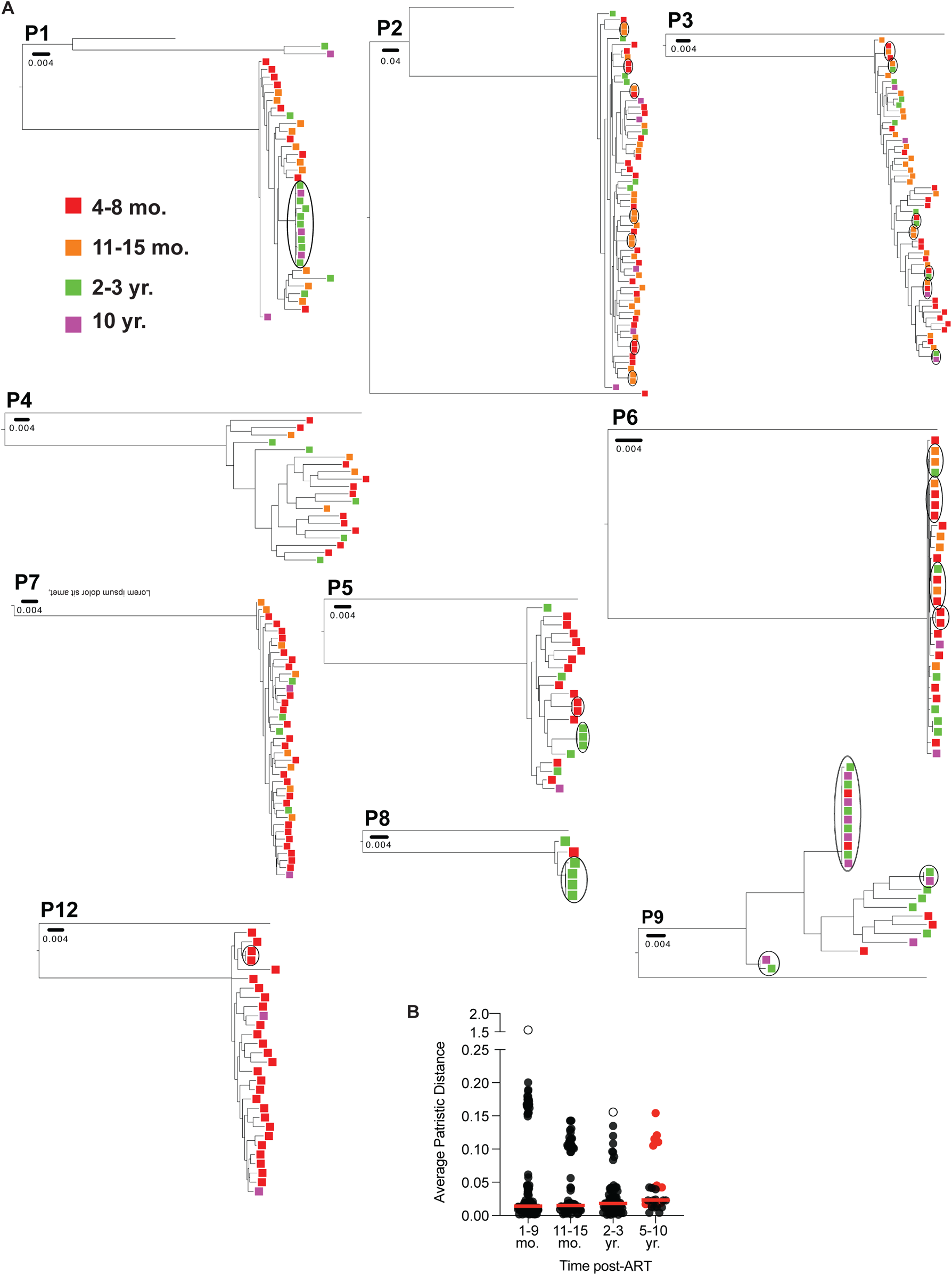
Diversity of the reservoir does not change over the course of long-term ART treatment. (A) Phylogenic tree of total HIV sequence of intact proviruses found in all participants (N=10). Branch lengths are proportion to a genetic distance (scale bar indicated below each Sample ID). Clones are outlined in black. Color of squares represents when the sequence was detected in the reservoir, as indicated by the key on the left. (B) Graph showing average patristic distance between each intact provirus and the rest of the population from the same time point of the same individual. Each dot represents one intact provirus found in the reservoir. Unfilled dots represent outliers that were excluded from analysis. Data derived from participants sampled at 10 years post-ART are highlighted in red. Horizontal red line indicates the mean patristic distance at each time point. Grubb’s test was used to calculate outliers, and A Kruskal-Wallis test with subsequent Dunn’s multiple comparisons was used to analyze data where appropriate.

**Table S2.** Definition of defective categories

| Defective Category | Characteristics of category |
| --- | --- |
| Missing Internal Genes | Truncated provirus, with large internal deletions spanning one or more coding genes |
| Non-coding | Full-length or truncated provirus, containing at least part of all genes, and early stop codons in one or more coding gene. |
| MSD Mutation | Full-length, in-frame, but lacking the 5' donor splice site 1 (D1) |
| Inversions or Duplications | Inversions or duplications detected in any genes or genomic sequence |
| Uncategorized | Unannotated defective due to missing either 3' LTR or 5' PS region where NFL primers should be binding |

**Supplementary Figure 3.**
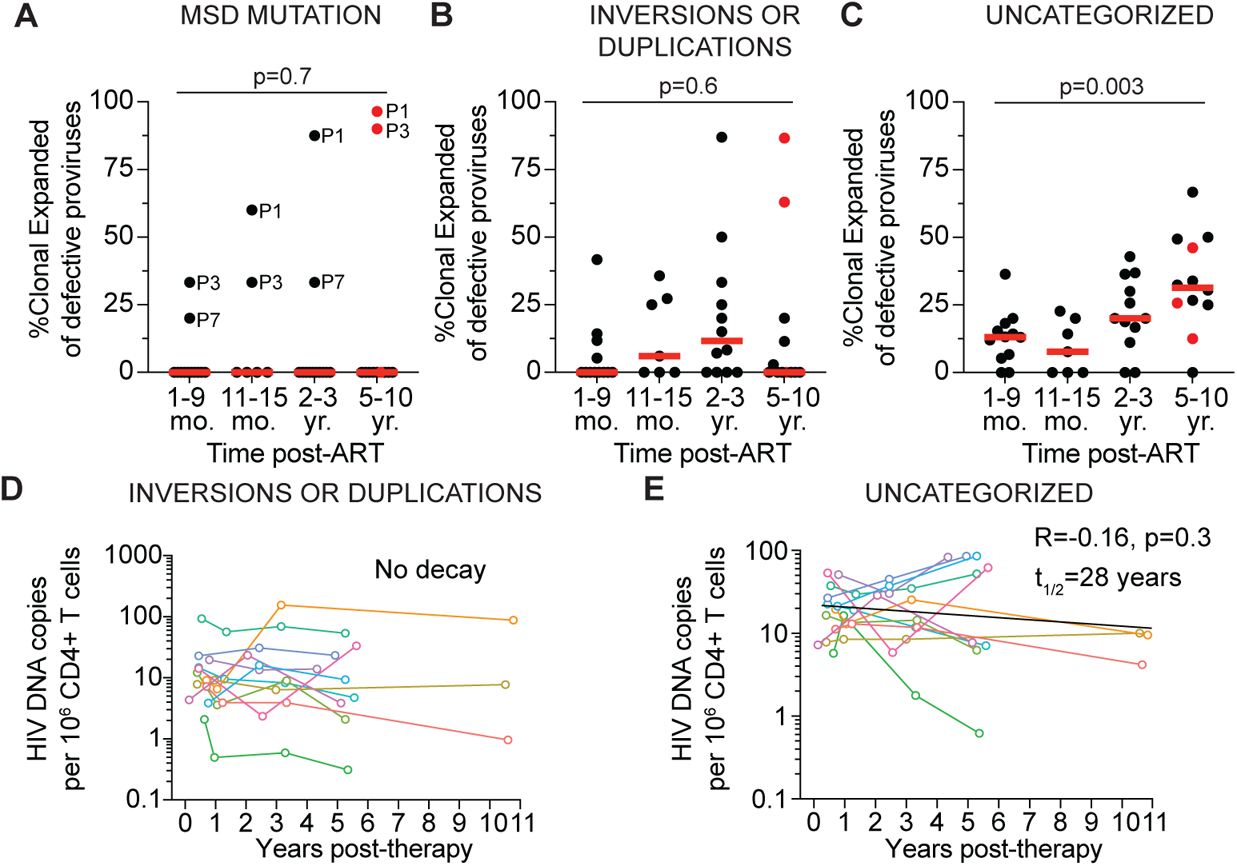
Longitudinal analysis of different categories of defective proviral reservoirs. Frequency of clonally expanded defective proviruses per individual for defective categorized as (A) MSD mutation (Sample identifiers added for the few samples with clonally expanded defectives categorized as MSD mutation), (B) Inversion or duplications, or (C) Uncategorized. Participants sampled at 10 years post-ART are highlighted in red. Red horizontal line indicated median frequency. Graph showing the number of defective HIV-1 proviruses per million CD4+ T cells detected over time, when defectives are categorized as (D) Inversions of duplications (No decay, no half-life calculated) or (E) Uncategorized (Half-life = 28 years, R=-0.16, p=0.3). Each colored line represents one individual participant. A Kruskal-Wallis test \with subsequent Dunn’s multiple comparisons was used to analyze data where appropriate. The exponential decay half-life was calculated by using log10 IUPM values and a linear regression model.

